# A Phosphopantetheinyl Transferase from *Dictyobacter vulcani* sp. W12 Expands the Combinatorial Biosynthetic Toolkit

**DOI:** 10.1101/2025.03.24.645039

**Authors:** Kenneth K. Hsu, Charlie M. Ferguson, Christina M. McBride, Nicholas B. Mostaghim, Kelsey N. Mabry, Robert Fairman, Yae In Cho, Louise K. Charkoudian

**Affiliations:** Department of Chemistry, Haverford College, Haverford, PA 19041; Department of Biology, Haverford College, Haverford, PA 19041

**Keywords:** phosphopantetheinyl transferase, acyl carrier protein, polyketide, synthase

## Abstract

The value of microbial natural product pathways extends beyond the chemicals they produce, as the enzymes they encode can be harnessed as biocatalysts. Microbial type II polyketide synthases (PKSs) are particularly noteworthy, as these enzyme assemblies produce complex polyaromatic pharmacophores. Combinatorial biosynthesis with type II PKSs has been described as a promising route for accessing never-before-seen bioactive molecules, but this potential is stymied in part by the lack of functionally compatible non-cognate proteins across type II PKS systems. Acyl carrier proteins (ACPs) are central to this challenge, as they shuttle reactive intermediates and malonyl building blocks between the other type II PKS domains during biosynthesis. Activating ACPs to their *holo* state via the phosphopantetheinyl transferase (PPTase)-catalyzed installation of a coenzyme A (CoA)-derived phosphopantetheine (Ppant) arm is critical to effectively study and strategically engineer type II PKSs, but not all ACPs can be activated using conventional PPTases. Here, we report the discovery of a previously unexplored non-actinobacterial PPTase from *Dictyobacter vulcani* sp. nov. W12 (vulcPPT). We explored its compatibility with both native and non-native ACPs, observing that vulcPPT activated all ACPs tested in this study, including a non-cognate, non-actinobacterial ACP which cannot be activated by the prototypical broad substrate PPTases AcpS and Sfp. Strategic optimization of phosphopantetheinylation reaction conditions increased *apo* to *holo* conversion. In addition to identifying a promising new promiscuous PPTase, this work establishes a road map for further investigation of PPTase compatibility and increases access to functional synthase components for use in combinatorial biosynthesis.

## INTRODUCTION

Microorganisms create a vast array of organic molecules that are not essential to the organism’s survival but confer them with an ecological competitive advantage. Type II polyketides are a particularly exciting class of these secondary metabolites because of their medicinally relevant properties (*e*.*g*. the antibiotic tetracycline and anticancer agent doxorubicin).^1^ These molecules are manufactured by multi-enzyme assemblies known as type II polyketide synthases (PKSs) which are spatially encoded within microbial genomes as biosynthetic gene clusters (BGCs).^1^ At minimum, a type II PKS comprises an acyl carrier protein (ACP), which must be post-translationally modified via the installation of a phosphopantetheine arm to its active “*holo*” form, and a ketosynthase-chain length factor (KS-CLF). The *holo*-ACP and KS-CLF interact to transform malonyl-based building blocks into a nascent beta-keto polyketide chain through a series of Claisen-like decarboxylation reactions.^1^ Subsequent reactions with a series of accessory enzymes tailor the polyketide intermediate into its final polyaromatic structure. The diversity of type II polyketides originates from variability in both the KS-CLF, which is a primary determinant in the number of carbons in the poly-beta-keto chain, and the members of the “tailoring enzyme roster” which catalyze the late-stage biochemical transformation and functionalization of the polyketide backbone.^1^ Unlike type I synthases (or synthetases), each protein or enzyme within a type II PKS exists as a discrete domain, making these systems uniquely conducive to mixing-and-matching synthase components in an effort to gain access to non-native chemical diversity. However, impaired ACP-protein interactions prevent such combinatorial biosynthesis efforts,^2^ so exploring ACPs that are more suitable for mediation of these interactions is crucial.

ACPs from non-actinobacterial type II PKSs are historically understudied and represent an interesting set of proteins to explore. While recent studies suggest that non-actinobacterial KS-CLFs are uniquely amenable to expression in *Escherichia coli*, their cognate ACPs can be expressed but not activated to their active *holo* form using conventional approaches.^3–6^ For example, the *Photorhabdus luminescens* TT01 type II PKS ACP could not be activated by the *E. coli* PPTase, AcpS, requiring the co-expression of two additional auxiliary enzymes for the successful *in vivo* production of the type II polyketide in *E. coli*.^4^ A similar barrier was encountered in the *in vitro* reconstitution of the *Gloeocapsa sp*. PC7428 type II PKS system in which neither Sfp nor AcpS could convert the *E. coli* heterologously expressed *apo-ACP* to its *holo* form, requiring strategic mutation of the ACP to enable type II PKS reconstitution *in vitro*.^3^

This activation, called phosphopantetheinylation, is catalyzed by an enzyme known as a phosphopantetheinyl transferase (PPTase) that installs a coenzyme A (CoA)-derived 18-Å long, 4’-phosphopantetheine prosthetic group (Ppant arm, Figure 1A) on a conserved serine located on the *N*-terminus of helix II of the ACP.^7^ The Ppant arm enables the ACP to tether and transport molecular building blocks and the growing polyketide chain between the other discrete type II PKS domains. It is well-documented that ACPs and peptidyl carrier proteins (PCPs) can be compatible with PPTases encoded by different BGCs within an organism;^8–10^ similarly, carrier proteins (CPs) from one organism can be activated by PPTases from different organisms altogether^11^. The PPTase from the *E. coli* fatty acid synthase (AcpS)^12^ and the PPTase from the *Bacillus subtilis* surfactin non-ribosomal peptide synthetase (Sfp)^13^ and its commonly used R4-4 variant^14^ (referred to as ‘Sfp’ from this point forward), are well-known for their promiscuity in converting non-cognate CPs to their *holo* form and are therefore widely used as biosynthetic tools. However, as noted above, these broad substrate PPTases have limited success with activating non-actinobacterial ACPs. Whilst the strategic mutation of non-actinobacterial ACPs can confer Sfp compatibility,^3^ identifying new PPTases which can readily activate wild type non-actinobacterial ACPs will improve access to type II PKS components. Herein, we report the *E. coli* heterologous expression and subsequent characterization of a previously unexplored PPTase encoded in the *Dictyobacter vulcani* sp. W12 genome (vulcPPT). vulcPPT demonstrates the ability to activate ACPs that are incompatible with Sfp and AcpS, and therefore represents an important new contribution to the biosynthetic toolkit.

**Figure 1.**
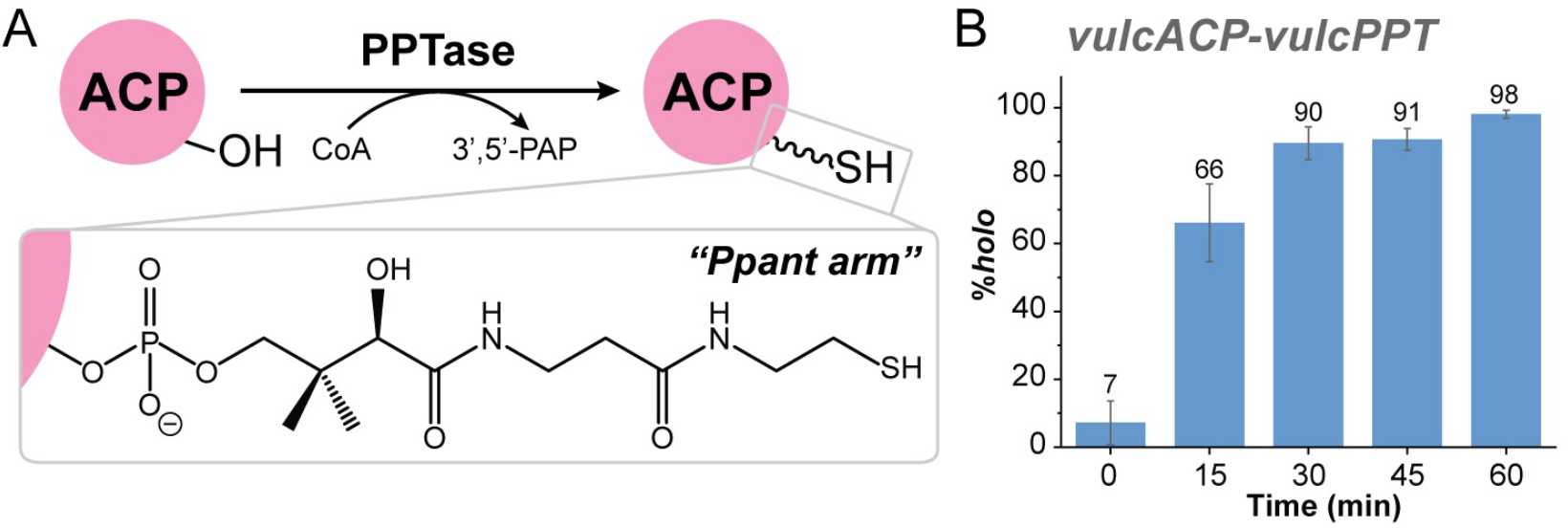
The PPTase from *Dictyobacter vulcani* sp. W12 (vulcPPT) converts its putative cognate ACP from the inactive *apo* form to active *holo* form. (A) ACP activation from *apo* to *holo* form is facilitated by a PPTase that installs the 18 Å CoA-derived phosphopantetheine (Ppant) arm on the conserved serine at the *N*-terminus of helix II. (B) The reaction of vulcACP:vulcPPT at over 0 to 60 min (see Supporting Information for reaction conditions), monitored by LC-MS, shows that vulcPPT efficiently activates its native ACP partner to nearly 100% within 60 min.

## RESULTS/DISCUSSION

To identify novel promiscuous PPTases that could readily activate non-actinobacterial ACPs, we turned to previously uncharacterized putative non-actinobacterial type II PKSs BGCs and their associated PPTases^15,16^ for *E. coli* heterologous expression and subsequent characterization. One such type II PKS BGC was identified in the *Dictyobacter vulcani* sp. W12 genome. Isolated from the soil of the Mt. Zao volcano in Japan, *D. vulcani* sp. W12 is a member of the *Ktedonobacteria* class, which is well known for actinomycete-like morphology and capability for secondary metabolite production.^17^ antiSMASH^18^ analysis of the *D. vulcani* sp. W12 genome revealed a putative type II PKS BGC with a unique, triad-like condensation (KS-CLF) domain architecture^16^ and an ACP with a non-canonical PPTase recognition motif (IDSI instead of LDSL). We selected the sole PPTase (vulcPPT) from the annotated protein list provided in the NCBI whole genome shotgun (WGS) Sequence Set Browser for *D. vulcani* sp. W12 for our heterologous expression efforts, but three additional proteins with homology to *Bacillus subtilis* Sfp and *Escherichia coli* AcpS were identified through BLASTp similarity searches. Interestingly, the genome is predicted to harbor >70 CPs across several putative non-ribosomal peptide synthetase (NRPS) and PKS BGCs, suggesting to us that vulcPPT might display unique and/or broad substrate activity.

The vulcPPT gene (see Methods and Table S1 in Supporting Information) was cloned into a pET28a-derived construct for *E. coli* heterologous expression with an *N*-terminal His_6_ tag and subsequently purified via affinity column purification to a yield of 60 mg/L. The protein was characterized via SDS-PAGE and liquid chromatography mass spectrometry (LC-MS; Supporting Figures S1 and S2, respectively). The AlphaFold 3^19^-predicted vulcPPT structure depicts vulcPPT as a pseudo-dimer consisting of two structurally similar subdomains attached by a polypeptide loop with high confidence (Figure S3). Together, the sequence, predicted structure, and size (249 aa) of vulcPPT suggest that vulcPPT is an Sfp-type PPTase.^7^ Circular dichroism (CD) wavelength experiments demonstrate that vulcPPT contains 26.8% α-helical, 8.4% antiparallel β-sheets, and 6.1% parallel β-sheets content.^20,21^ Further protein melting experiments reveal a melting temperature (T_melt_) of 45.53 °C (± 0.15 °C) for vulcPPT (Figure S4 and Table S2). Preliminary examination of its *in vitro* activity demonstrates that vulcPPT can fully convert its putative cognate ACP (hereafter referred to as vulcACP) from the inactive *apo* form to active *holo* form within an hour (Figure 1B and Supporting Information).

To evaluate vulcPPT’s substrate scope relative to PPTases routinely used in the field, we assessed the ability of vulcPPT, AcpS, and Sfp to phosphopantetheinylate four ACPs: (i) vulcACP, (ii) the ACP from the *E. coli* fatty acid synthase (FAS) system (AcpP), (iii) the ACP from the *Streptomyces coelicolor* actinorhodin type II PKS (actACP), and (iv) a previously unexplored ACP from the *Zooshikella* sp. WH53 putative type II PKS BGC (zooACP). These four ACPs were selected to represent diverse substrates, including the putative cognate ACP for vulcPPT (i), the prototypical FAS ACP and native substrate for AcpS (ii), the prototypical type II PKS ACP (iii), and a never-before-studied non-cognate non-actinobacterial ACP (iv). Plasmids encoding for the four ACPs were transformed into *E. coli* BL21 (DE3) cells for expression in their majority (∼90– 95%) inactive *apo* form. Phosphopantetheinylation reactions were performed by reacting an *apo*-ACP with a PPTase in the presence of CoA, dithiothreitol (DTT), and MgCl_2_ before being quenched at 18 hours with formic acid and analyzed via liquid chromatography mass spectrometry (LC-MS). Under the LC conditions used (see Supporting Information for details) the two forms of the ACP (*apo* and *holo*) elute at distinct retention times, allowing for percent conversion to be quantified.

VulcPPT fully converts *apo*-actACP and *apo*-AcpP to their active *holo* states, matching the activity of both AcpS and Sfp (Figure 2B). However, vulcPPT was significantly more efficient in converting its cognate *apo*-vulcACP to its *holo* form than AcpS (100% (± 0.0%) versus 58% (± 4.5%), respectively). Interestingly, of the three PPTases studied, only vulcPPT was capable of converting zooACP to its *holo* state, albeit at low efficiency (∼11% (± 1.5%) *holo*, Figure 2B) under non-optimized conditions. Together these data highlight the broad substrate scope of vulcPPT and its utility in obtaining *holo*-ACPs that cannot be readily obtained using field-standard PPTases. These results are particularly significant in the context of recent work on cyanobacterial PPTases and ACPs, in which diverse PPTases could not outperform Sfp in activating cyanobacterial CPs.^22^

**Figure 2.**
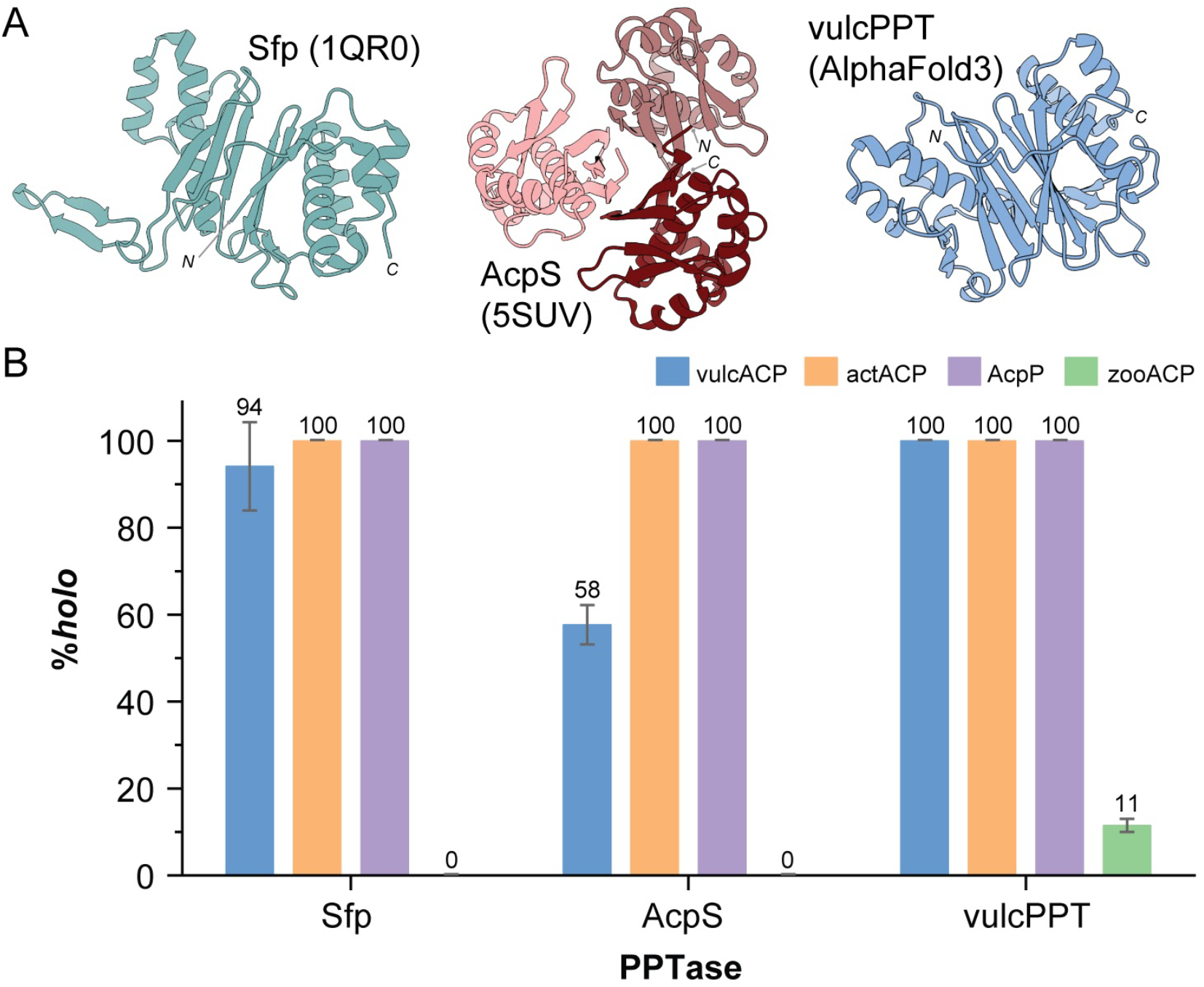
The PPTase from *Dictyobacter vulcani* sp. W12 (vulcPPT) shows expanded substrate scope compared to Sfp and AcpS for the ACPs studied. (A) The structures of Sfp (PDB 1QR0), AcpS (PDB 5SUV), and vulcPPT (AlphaFold3 prediction) suggest that vulcPPT belongs to the Sfp-family of PPTases. (B) VulcPPT can activate a range of ACPs, including ACPs that are not readily activated using AcpS and Sfp. When *apo*-vulcACP (blue), *apo*-actACP (orange), *apo*-AcpP (purple), and *apo*-zooACP (green) were reacted with Sfp, AcpS, and vulcPPT (see Supporting Information for reaction conditions) for 18 hrs, vulcACP, actACP, AcpP were converted to their *holo* forms with efficiencies ranging from 58% (± 4.5%) to 100% (± 0.0%) by all three PPTases. In contrast, zooACP was only activated by vulcPPT, achieving a conversion rate of 11% (± 1.5%). See Figures S5-S8 for corresponding LCMS data.

Given promising preliminary results, we next sought to determine whether the *apo* to *holo* conversion efficiency observed for the zooACP-vulcPPT reaction could be improved by altering the phosphopantetheinylation conditions. The four reaction variables--(i) temperature, (ii) pH, (iii) vulcPPT concentration, and (iv) CoA concentration--were assessed independently with 25 °C, pH 7.6, 1.0 μM vulcPPT, and 10x CoA as the standard conditions. First, a 35 °C incubation temperature yielded the highest conversion to *holo-*zooACP (85% (± 2.2%)) within the range 20– 37 °C (Figure 3A). Second, vulcPPT produced the highest % *holo* conversion (27% (± 1.9%)) at pH 7.5, with a broad pH range of activity (Figure 3B). Interestingly, optimal conversion from *apo*-zooACP to *holo*-zooACP was observed at 3.0 μM vulcPPT and 750 μM CoA (Figures S11 and S12, respectively). Taken together, these experiments reveal that the vulcPPT phosphopantetheinylation reaction conditions can be tuned to improve conversion efficiency.

**Figure 3.**
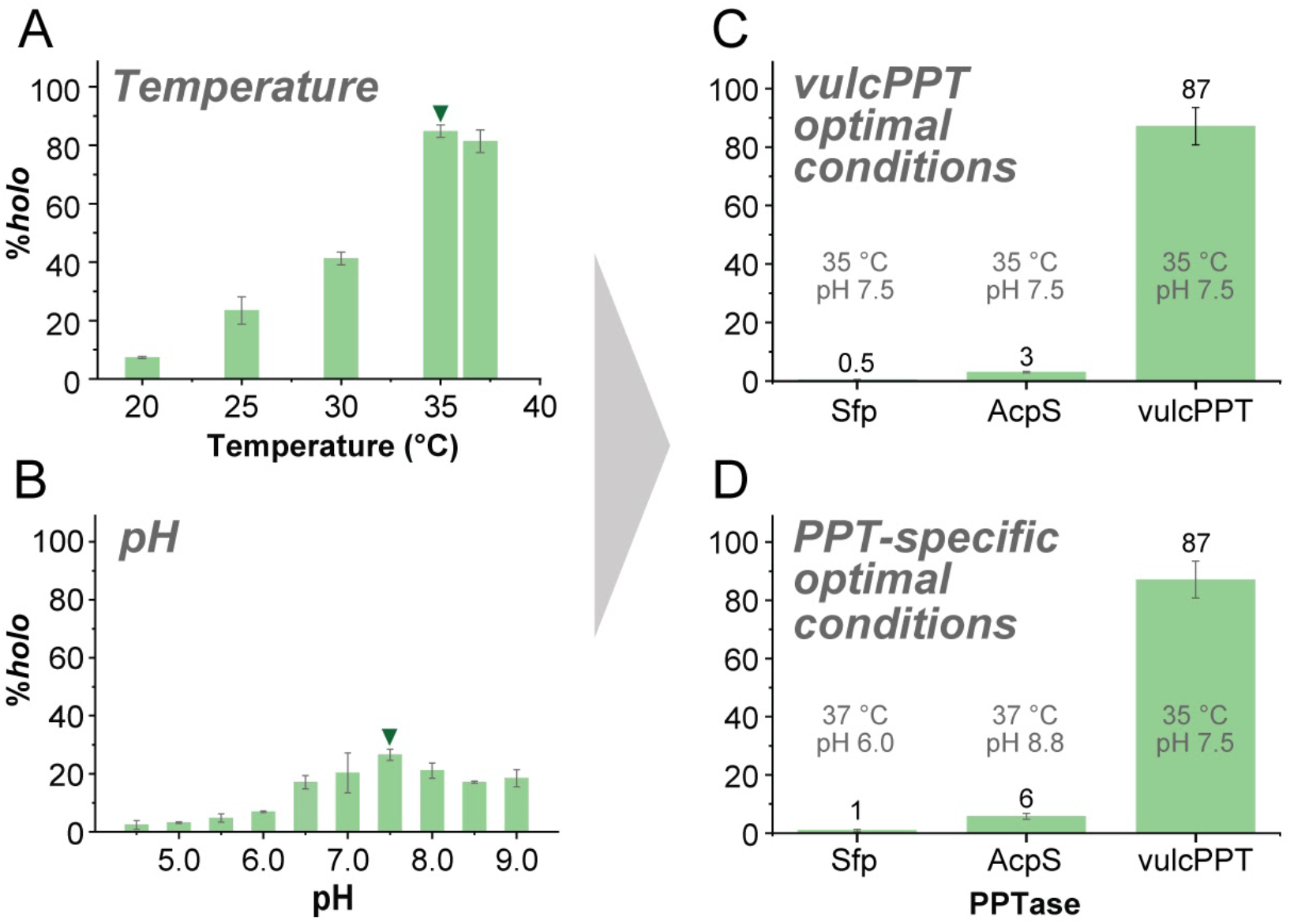
By optimizing reaction conditions, the phosphopantetheinylation of zooACP by vulcPPT could be improved from 11% (± 1.5%) to 87% (± 6.4%). To optimize the activation of *apo*-zooACP by vulcPPT, different variables were tested: temperature at a constant pH of 7.6 (A, more detailed version in Figure S9), pH at a constant temperature of 25 °C (B, more detailed version in Figure S10), vulcPPT concentration (Figure S11), and CoA concentration (Figure S12). The condition with the highest percent conversion for each condition explored is marked with a triangle. (C) Under the optimized conditions for vulcPPT [35 °C, 750 μM (5 molar equivalents relative to *apo*-zooACP = 5x) CoA, 3.0 μM PPT, pH 7.5], *apo*-zooACP was converted to *holo*-zooACP at 87% (± 6.4%), which is 8-fold higher than the initial conditions explored. (D) Phosphopantetheinylation of *apo*-zooACP by Sfp, AcpS, and vulcPPT under their respective optimal conditions—pH 6.0 at 37 °C for Sfp, pH 8.8 at 37 °C for AcpS, and pH 7.5 at 35 °C for vulcPPT—followed by analysis after 18 hours. The results confirm that only vulcPPT effectively activates zooACP into its *holo* form under these conditions.

Next, vulcPPT phosphopantetheinylation reactions were performed on *apo*-zooACP using the conditions determined to be optimal for each variable explored (3.0 μM vulcPPT, 35 °C, pH 7.5, and 750 μM (5x) CoA, Figure 3C). Under these conditions, we observed over 87% (± 6.4%) conversion of *apo*-zooACP to its *holo* form. The nearly 8-fold improvement in efficiency compared to the result from unoptimized conditions (11% (± 1.5%)) seems to be primarily influenced by temperature (Figure 3A). The zooACP phosphopantetheinylation reaction was performed under identical conditions using Sfp and AcpS in place of vulcPPT which yielded only 0.5% (± 0.2%) and 3% (± 0.3%) *holo*-zooACP, respectively (Figure 3C). Finally, to more accurately compare the efficiency of all three PPTases, phosphopantetheinylation reactions were conducted on *apo*-zooACP using the literature-reported optimal conditions for Sfp (pH 6.0, 37 °C, Figure 3D) and AcpS (pH 8.8, 37 °C, Figure 3D).^23,24^ These reactions yielded 1% (± 0.4%) and 6% (± 1.0%) *holo*-zooACP, respectively–an improvement over the initial reactions using vulcPPT optimal conditions (pH 7.6, 35 °C), yet still much lower than the >87% *holo-*zooACP achieved by vulcPPT. These data support the conclusion that among the three PPTases studied, vulcPPT is the most effective at activating zooACP. Compared to the optimal pH and temperature of AcpS and Sfp, vulcPPT behaves optimally at less harsh pH conditions (pH 7.6) and cooler temperatures (35°C). VulcPPT is therefore useful for reactions where substrates require more amenable conditions. We hypothesized that the broad substrate compatibility of vulcPPT is derived from its ability to recognize a broader spectrum of ACP motifs, as compared to Sfp and AcpS.^25^ To gain further insight, the *D. vulcani* sp. W12 genome was analyzed using antiSMASH^18^ to collate a set of putative CPs, including ACPs from type I/II PKSs and fatty acid synthases as well as PCPs from non-ribosomal peptide synthetases. 79 CP genes were identified and further analyzed via multiple sequence alignment (Figure S13). Notably, only a small percentage (6.3%) of these CPs contained the traditionally Sfp-favored amino acid motif of DSL (where the S is the conserved serine at the *N*-terminus of helix II that is the attachment point for the Ppant arm). Instead, this DSL motif was frequently replaced by other motifs, such as DSI (44.3%, 35/79) and HSL (34.1%, 27/79) among others. These data suggest that vulcPPT, which is the sole PPTase identified from the annotated protein list in the NCBI whole genome shotgun (WGS) Sequence Set Browser for *D. vulcani* sp. W12, may be capable of recognizing a broader range of interaction motifs. This sort of “crosstalk”, where a PPTase activates more than one ACP in an organism (as opposed to having each PPTase be specific to a single CP), has been observed in recent years and can be leveraged in strategic engineering work.^8–11,26^ Additional evidence to support this hypothesis lies in the fact that a DST motif has been identified as a key residue that makes an ACP incompatible with Sfp.^3^ Moreover, basal expression of the *E. coli* AcpS could not convert the *Rhizobia* CP SMb20651, which contains a DST motif, to its *holo* form. Conversion was only achieved by co-overexpression of the *E. coli* or *S. meliloti* AcpS, suggesting that the presence of a DST motif instead of a DSL motif presents barriers to accessing CPs in their *holo* form.^7^ ZooACP features a DST motif and cannot be converted by Sfp to its *holo* form in any meaningful quantity. In comparison, vulcPPT has demonstrated the ability to convert zooACP to its *holo* form in significant quantities, suggesting that vulcPPT is compatible not only with the more accessible DSL, DSI, and HSL motifs but also with the previously difficult to access DST motif.

The impact of discovering and characterizing new PPTases is highlighted by the ability of vulcPPT to convert *apo*-zooACP, a non-cognate CP which was previously inaccessible using conventional PPTases, to its active *holo* form. It is critical to expand the biosynthetic toolkit such that diverse *holo*-ACPs can be accessed given that 1) *holo*-ACPs are an essential component to any PKS, whether native or created via combinatorial biosynthesis, and 2) impaired ACP-protein interactions lead to failure of a PKS to produce a polyketide product. PPTases like vulcPPT allow us to obtain ACPs that are not accessible in their WT form and can currently only be activated by Sfp if the ACP is strategically engineered.^3,26^ We hope that the introduction of vulcPPT to the combinatorial biosynthetic toolkit will improve access to functional ACPs while also providing additional clues that can be used to uncover the molecular underpinnings of PPTase-ACP compatibility.

## ASSOCIATED CONTENT

### Supporting Information

The Supporting Information is available free of charge at [doi link will be inserted here later].

Detailed descriptions of materials and methods; plasmid, primer, and amino acid sequence information; SDS-PAGE of the purified ACPs and phosphopantetheinyl transferases; LC-MS spectra of the *apo-* and *holo-*ACPs upon phosphopantetheinylation; CD spectrum and T_melt_ of vulcPPT; temperature, pH, vulcPPT concentration, and coenzyme A concentration optimization of the phosphopantetheinylation of *apo*-zooACP by vulcPPT; multiple sequence alignment of 79 CPs encoded by the *Dictyobacter vulcani* sp. W12 genome.

## Author Contributions

The manuscript was written through contributions of all authors. All authors have given approval to the final version of the manuscript. *Co-corresponding authors.

C.M.M., C.M.F., K.K.H., L.K.C., N.B.M., K.N.M., R.F., Y.I.C. designed the research. C.M.M., C.M.F., K.K.H., N.B.M., K.N.M., Y.I.C. collected and analyzed data. C.M.M., C.M.F., K.K.H., L.K.C., Y.I.C. wrote the manuscript.

## Funding Sources

We are grateful for generous support from the National Science Foundation (CHE2201984 to L.K.C.), a 2021 Arnold and Mabel Beckman Foundation Scholarship and 2022 Goldwater Scholarship (C.M.M) and Haverford College.

## ACKNOWLEDGMENTS

We thank Professor Joris Beld (Drexel University) for helpful discussions as well as Gabriel Rocco Sotero for technical support. We are also grateful to the 2023 Haverford College Laboratory in Biochemical Research class for their support and guidance.

## ABBREVIATIONS

PKS, polyketide synthase; BGC, biosynthetic gene cluster; ACP, acyl carrier protein; PPTase, phosphopantetheinyl transferase; CoA, coenzyme A; AcpS, *holo*-(acyl carrier protein) synthase; Sfp, 4’-phosphopantetheinyl transferase protein from *Bacillus subtilis*; NRPS, nonribosomal peptide synthetase; KS-CLF, ketosynthase chain length factor; Ppant arm, 4’-phosphopantetheine prosthetic group; SDS-PAGE, sodium dodecyl sulfate polyacrylamide gel electrophoresis; LC-MS, liquid chromatography mass spectrometry; CD, circular dichroism.

## TABLE OF CONTENTS GRAPHIC

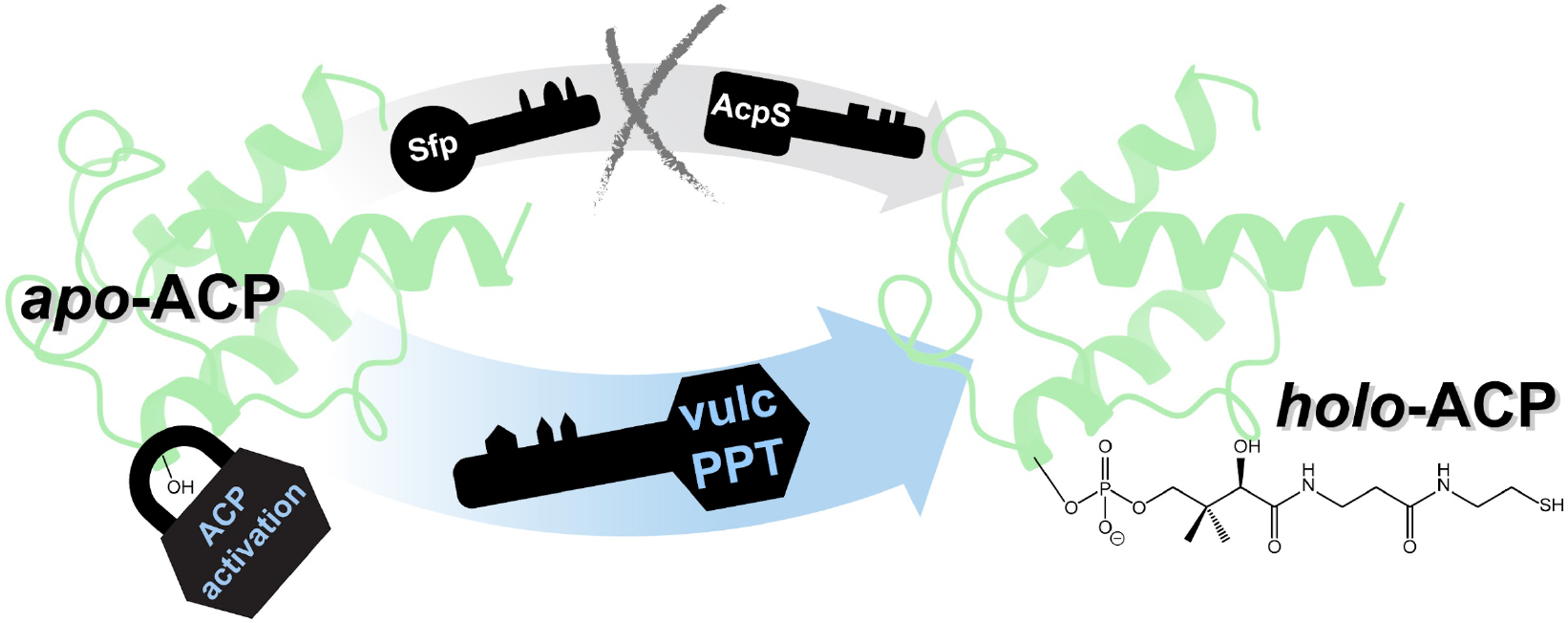

## Notes

### Competing Interest Statement

The authors have declared no competing interest.

