## Supporting Information for "A Phosphopantetheinyl Transferase from *Dictyobacter vulcani* sp. W12 Expands the Combinatorial Biosynthetic Toolkit"

| **Table of Contents** | **Page** |
|---|---|
| Materials and Methods | 3–5 |
| **Table S1.** Plasmid, primer, and amino acid sequence of all proteins used in this study. | 6 |
| **Table S2**. Thermal stability curve fit of vulcPPT. | 7 |
| **Figure S1.** SDS-PAGE characterization of the expressed and purified acyl carrier proteins (ACPs) and phosphopantetheinyl transferases (PPTases) used in this study. | 8 |
| **Figure S2.** LC-MS spectrum of purified vulcPPT. | 9 |
| **Figure S3.** Predicted structure of vulcPPT (AlphaFold 3) and its overlay with Sfp. | 10 |
| **Figure S4.** Circular dichroism spectrum and $T_{melt}$ of vulcPPT. | 11 |
| **Figure S5.** Mass spectra of *apo*- and *holo*-vulcACP, and LC chromatograms after PPTases reactions with Sfp, AcpS and vulcPPT. | 12 |
| **Figure S6.** Mass spectra of *apo*- and *holo*-actACP, and LC chromatograms after PPTases reactions with Sfp, AcpS and vulcPPT. | 13 |
| **Figure S7.** Mass spectra of *apo*- and *holo*-AcpP, and LC chromatograms after PPTases reactions with Sfp, AcpS and vulcPPT. | 14 |
| **Figure S8.** Mass spectra of *apo*- and *holo*-zooACP, and LC chromatograms after PPTases reactions with Sfp, AcpS and vulcPPT. | 15 |
| **Figure S9.** Temperature optimization of the phosphopantetheinylation of *apo*-zooACP by vulcPPT (detailed version of Fig 3A). | 16 |
| **Figure S10.** pH optimization of the phosphopantetheinylation of *apo*-zooACP by vulcPPT (detailed version of Fig 3B). | 16 |
| **Figure S11.** VulcPPT concentration optimization of the phosphopantetheinylation of *apo*-zooACP by vulcPPT. | 17 |
| **Figure S12.** Coenzyme A concentration optimization of the phosphopantetheinylation of *apo*-zooACP by vulcPPT. | 17 |
| **Figure S13.** Multiple sequence alignment of 79 carrier proteins encoded by the *Dictyobacter vulcani* sp. W12 genome. | 18 |
| References for Supporting Information | 19 |

### Materials and Methods

**Molecular Cloning of vulcPPT, vulcACP, and zooACP.**
The plasmids encoding for vulcPPT, vulcACP, and zooACP with *N*-terminal His-tags were created as part of a course-based undergraduate research experience (CURE) in 2023. Template DNAs were purchased from Twist Bioscience, and the primers (both forward and reverse) were purchased from Eurofins Genomics (**Table S1**). Each template DNA was amplified via polymerase chain reaction (PCR) under the following conditions: 22.5 µL nuclease free water, 10.0 µL 5X Q5 GC enhancer, 10.0 µL 5X Q5 reaction buffer, 2.5 µL forward primer, 2.5 µL reverse primer, 1.0 µL dNTPs, 1.0 µL template DNA for a total volume of 50 µL. In a Bio-Rad S1000™ Thermocycler, the reaction mixture underwent a seven-step cycling procedure as follows: 1. 98 °C for 30 sec, 2. 98 °C for 10 sec, 3. 72 °C for 30 sec, 4. 72 °C for 30 sec, 5. go to step 2, 29 times, 6. 72 °C for 2 min, 7. hold at 4 °C. Amplification of target DNA was verified via DNA gel electrophoresis (1% (w/v) agarose) and the amplified product was removed and purified using the Zymoclean™ Gel DNA Recovery Kit. The pET28a vector was linearized through digestion with NdeI and EcoRI and purified via DNA gel electrophoresis followed by gel extraction (Zymoclean).

Plasmids were constructed via Gibson Assembly[1] by inserting the sequence of interest into a pET28a vector at the NdeI and EcoRI cut sites. Plasmid inserts and vectors were combined with Gibson Assembly® Master Mix (New England BioLabs), consisting of T5 Exonuclease, Phusion Polymerase, Taq Ligase, dNTPs, and $MgCl_2$ in Tris-HCl buffer. The reaction mixture was incubated at 50 °C on a heat block. Plasmids encoding for the expression of the actinorhodin ACP (actACP; MC002067), Sfp R4-4, and AcpP were generously provided by the Chang Lab (Princeton University), Lin Lab (Georgia State University), and Khosla Lab (Stanford University), respectively. For details of the plasmids, see **Table S1**.

**Expression and Purification of ACPs/PPTases.**
Plasmids were transformed into competent BL21 (DE3) cells for expression. Seed cultures (10 mL LB supplemented with kanamycin (kan) at 50 µg/mL) were grown at 37 °C and added to production cultures (1 L LB with 50 µg/mL kan). Production cultures were incubated with gentle shaking at 37 °C and induced with isopropyl β-D-1-thiogalactopyranoside (IPTG; 250 µM final concentration) at $OD_{600}$ 0.4–0.6. Cells were harvested by centrifugation (17,000 × *g*, 4 °C, 20 min per round), resuspended in lysis buffer (50 mM sodium phosphate buffer, 10 mM imidazole, 300 mM NaCl, pH 7.6) and sonicated (40% amplitude, 30 sec cycles, 10 min) on ice. Lysed cell suspensions were centrifuged (17,000 × *g*, 4 °C, 1 hr) and the supernatant was mixed with Ni-NTA agarose beads (Gold Biotechnology) at 4 °C for 1.5–2 hrs with gentle rocking. The mixture was applied to a poly-prep column, and washed with 5 column volume (CV) of lysis buffer, followed by 50 mL of wash buffer (50 mM sodium phosphate, 30 mM imidazole, 300 mM NaCl, pH 7.6) before being eluted with 10 mL of elution buffer (50 mM sodium phosphate, 250 mM imidazole, 100 mM NaCl, 10% (v/v) glycerol, pH 7.6). Ten 1-mL elution fractions were obtained and fractions with A280 values > 0.15 were pooled together and dialyzed against a storage buffer (50 mM sodium phosphate, 10% (v/v) glycerol, pH 7.6) overnight. Protein concentrations were determined by Nanodrop 2000 spectrophotometer (Thermo Fisher Scientific). Proteins were flash frozen, and ACPs were stored at 205–437 µM and PPTases were stored at 67–370 µM. Proteins were characterized by SDS-PAGE and LC-MS (see below for details).

**Method to Quantify *apo*-ACP to *holo*-ACP Conversion.**
Reactions converting *apo*-ACP to *holo*-ACP were routinely run in triplicate using the following protocol unless otherwise specified. For a 125 µL-reaction, *apo*-ACP was incubated with the PPTase in a reaction mixture using stock solutions of 50 mM dithiothreitol (DTT), 50 mM coenzyme A (CoA, lithium salt from CoALA Biosciences), and 1 M $MgCl_2$ in 50 mM sodium phosphate buffer, pH 7.6 at 25 °C (final concentrations of each reaction component: 150 µM *apo*-ACP; 1 µM PPTase; 2.5 mM DTT; 1.5 mM CoA; 10 mM $MgCl_2$).
To adjust the pH of each reaction mixture, the *apo*-ACP stock in 50 mM sodium phosphate buffer (pH 7.6) was concentrated to 750 µM such that the ACP occupied only 20% of the final reaction volume. The remaining reaction volume was reached by adding 50 mM sodium phosphate buffer of varying pH. To stop the reaction, 50 µL of the reaction mixture was removed and quenched with 10 µL of 25% formic acid for 30 minutes. The quenched sample was prepared for LC-MS by adding 20 µL of 5% NaOH and 20 µL of LC-MS grade water. For more detailed LC-MS methods and the percent quantification of *apo/holo* of these ACP samples, see below.

**Liquid Chromatography-Mass Spectrometry (LC-MS).**
An Agilent Technologies InfinityLab G6125B LC/MS coupled with an Agilent 1260 Infinity II LC system equipped with a Waters XBridge Protein BEH C4 reverse phase column (300 Å, 3.55 µm, 2.1 mm × 50 mm) and XBridge Protein BEH C4 Sentry guard cartridge (300 Å, 3.5 µm, 2.1 mm × 10 mm) was used to analyze samples at 45 °C by electrospray ionization mass spectrometry (ESI MS) in positive mode. The instrument was equipped with two LC-MS grade solvents: Solvent A ($H_2O$ + 0.1% formic acid) and Solvent B (acetonitrile + 0.1% formic acid). The following gradients were used depending on the sample:

- **For the samples not requiring the *apo/holo* quantification:** 0–1 min 5% B; 1–3.1 min 95% B; 3.1–4.52 min 95% B; 4.52–4.92 min 5% B; 4.92–9 min 5% B. Unless otherwise stated, LC-MS samples were prepared by diluting 10 µL of protein sample with 90 µL of LC-MS grade water to the approximate concentration of 5–10 µM. Samples were injected (20 µL) and run using a capillary voltage of 3000 V and a fragmentation voltage of 75 V.
- **For the ACP samples to confirm the *apo/holo* quantification:** The C4 column was first equilibrated for 10 min at a 0.400 mL/min flow rate with 10% B, after which 20 µL of sample was injected into the column. The following solvent gradient was used post injection: 0–5 min 30% B; 5–30 min 50% B; 30–36 min 95% B; 36–41 min 10% B. Samples were run using a capillary voltage of 3000 V and a fragmentation voltage of 75 V.

Acquired mass spectra were deconvoluted using ESIprot online[2] and the observed and calculated molecular weights (MWs) were compared to confirm a successful phosphopantetheinylation. Absorbance data from the liquid chromatography instrument at 280 nm were zeroed and plotted in Origin 2023b.[3] Relative quantities of *apo*- and *holo*-ACP were calculated by integrating the respective UVvis absorbance of eluent peaks.

**Sodium Dodecyl Sulfate-Polyacrylamide Gel Electrophoresis (SDS-PAGE).**
For non-reducing SDS-PAGE samples, 5 µL of 6x purple gel loading dye (New England Biolabs) were combined with 25 µL protein samples (20 µM for ACPs; 15 µM for PPTases). For reducing SDS-PAGE samples, 5 µL of 6x purple gel loading dye and 1.5 µL of β-mercaptoethanol were combined with 23.5 µL protein samples (20 µM for ACPs; 15 µM for PPTases), making 5% (v/v) β-mercaptoethanol overall. Samples were denatured at 100 °C for 5 min and 10 µL of each sample was loaded in each well of the gel (Bio-Rad Mini-PROTEAN® TGX™ 10-well, 30 µL 4–20% precast polyacrylamide gels). SDS-PAGE gels were run in 1X

SDS-PAGE running buffer (from the 10X: 0.25 M Tris base, 1.92 M glycine, 1% (w/v) SDS, pH 8.3) at 120 V. Gels were washed with $ddH_2O$, stained with ThermFisher GelCode$^{TM}$ Blue for 1 hour with gentle shaking, destained overnight with $ddH_2O$ with gentle shaking, and imaged using a Bio-Techne FluorChem M System (**Figure S1**).

**Circular Dichroism.**
CD spectra were collected using an Aviv Model 410A circular dichroism spectropolarimeter. Protein samples were diluted to 200 µL to final concentrations of 10 µM in the storage buffer (50 mM sodium phosphate, 10% (v/v) glycerol, pH 7.6). Samples were injected into a High Precision Quartz SUPRSIL cuvette with 0.1 cm pathlength (Hellma Analytics). The spectropolarimeter was purged with nitrogen for two hours. After the UV lamp was warmed up for 30 minutes, CD spectra were collected at 25 °C with a range of 190–250 nm using bandwidth of 1 nm, a 0.5 nm step size, averaging time of 3 seconds, and 5 scans. To assess thermal stability, changes in signal at 222 nm were monitored as a function of temperature under the following parameters: 10–90 °C, 2-min equilibration, heating rate 2 °C $min^{-1}$, 30 sec signal averaging time, and 1 nm bandwidth. Pre- and post-melting spectra were smoothed using a smoothing function implemented in the Aviv software, using a window width of 11 data points, degree 2. The resulting spectrum (**Figure S4**) was converted to units of mean residue ellipticity (MRE) using the protein's amino acid sequence and the sample concentration. Analysis of protein secondary structure characteristics was conducted by uploading normalized data in units of MRE to the web server BeStSel.[3,4] Normalized data was plotted in Origin 2023b. To calculate the melting temperature of a protein ($T_{melt}$), changes in signal at 222 nm as a function of temperature were plotted in Origin 2023b. A sigmoidal curve was fitted to the plot using a Levenberg-Marquardt algorithm (logistic function) and the E50/x0 value was taken as the $T_{melt}$ of vulcPPT (**Figure S4 and Table S2**).[5]

### Tables and Figures

**Table S1**. Plasmid, primer, and amino acid sequence of all proteins used in this study.

| | |
|---|---|
| vulcPPT | Forward Primer:<br>CCT GGT GCC GCG CGG CAG CCA TAT GAT CGA AGA CAT CTG GC<br>Reverse Primer:<br>AAG CTT GTC GAC GGA GCT CGA ATT CTC ACA ACG TGC CGT TCC ATT GC<br><br>Insert:<br>ATGATCGAAGACATCTGGCAGCCACCTCCCTCAACATTAAAACTTGAGCAAAGCGCAGTCCAC<br>GTGTGGCGTGCTGACCTTCGTGCCTCTCAAGAATCGGTCGAACGTTTCCGCCACATCCTTTCGC<br>CTGAGGAGCAGGCACGCGCGCAGCGTTTCTACTTCGAACGTGATCGTTACCGTTGGACTATTGC<br>ACATGGCATCTTGCGTATCCTTTTGGCCCGTTACACTGGGCAAGATCCACGCGCCTTGCGCTTT<br>CAAGTCAACGCCTACGGGAAGCCTTCCCTGGTGCAGCCCGATCAACAACCCCGCCTGGAATTT<br>AACCTTAGCCACTCTCATGAGATGGCTTTATATGCTTTTACGTGGCAGCGCCAGATTGGGGTGG<br>ACGTAGAGTACATGCGTGATGATATTGGGTATGAAGAATTGGCTCGCCACTCCTTCAGCCCGA<br>CCGAGCAAGCCGTTCTTCTGTCGTTGGCGACCTCGCAACAAAAAGCCGCTTTCTTTAAATGCTG<br>GAGTTCCAAGGAAGCTTATATCAAGGGTCGCGGGATGGGTCTGTCTTTAGAGCTTAACCTTTTT<br>GACGTGGCTCTGGCTCCCGACAAGCCGGTTGCACTGTTAGCTTCCCGTGAAGACCCAGCCGAA<br>GTCCAACGCTGGTCAATGGCAAAACTGGAACCGGGTGCCGACTACGCTGGTGCGTTGGCGGTC<br>GAAGGGCCTCTTCCGGATATCTCTTGTTGGCAATGGAACGGCACGTTGTGA<br><br>Amino Acid Sequence:<br>MGSSHHHHHHSSGLVPRGSHMIEDIWQPPPSTLKLEQSAVHVWRADLRASQESVERFRHILSPEEQ<br>ARAQRFYFERDRYRWTIAHGILRILLARYTGQDPRALRFQVNAYGKPSLVQPDQQPRLEFNLSHSH<br>EMALYAFTWQRQIGVDVEYMRDDIGYEELARHSFSPTEQAVLLSLATSQQKAAFFKCWSSKEAYI<br>KGRGMGLSLELNLFDVALAPDKPVALLASREDPAEVQRWSMAKLEPGADYAGALAVEGPLPDISC<br>WQWNGTL |
| vulcACP | Forward Primer:<br>CCT GGT GCC GCG CGG CAG CCA TAT GGT CAA TGA CGC TCT GC<br>Reverse Primer:<br>AAG CTT GTC GAC GGA GCT CGA ATT CTC ACT TGC TGG CGG CTT CC<br><br>Insert:<br>ATGGTCAATGACGCTCTGCAAGAGACGATTTATCCGCGCGTAGTAGCTATTTTACGTTGCCAAG<br>TAGACGAAGACGATGAAATTCGCTCTGATACCGACCTGTTAACAGATTTAGGAATTGACTCAA<br>TCGGCCAAGTCGAAATCTGCCTTGCGCTGGAGAAAGAGTTCGGACTGCGCTTCTCCATTGCAG<br>AGCTTCGCGTTTGCACGACAGTAGATGAGGTGGTTCAGCTTGTCGTACACACGATTGCGGGTA<br>AGGAAGCCGCCAGCAAGTGA<br><br>Amino Acid Sequence:<br>MGSSHHHHHHSSGLVPRGSHMVNDALQETIYPRVVAILRCQVDEDDEIRSDTDLLTDLGIDSIGQVE<br>ICLALEKEFGLRFSIAELRVCTTVDEVVQLVVHTIAGKEAASK |
| zooACP | Forward Primer:<br>CCT GGT GCC GCG CGG CAG CCA TAT GAC CAA CGA AAC GGT TA<br>Reverse Primer:<br>AAG CTT GTC GAC GGA GCT CGA ATT CCT AAG CGG CCT CGC TTA AA<br><br>Insert:<br>ATGACCAACGAAACGGTTAAGGCCATCTACACAACGCTGGCGGAGTATTTGGATATG<br>CCGGTGTCTGAGTTGAAGGAGAATACAAACTTGGAAAACGAACTTCTGTTGGACTCT<br>ACTGAATTGGTCTGCATTATGGTAGCCTTGGAAAAATCTATGGATATTAGTCTGAAA<br>AATGTGCCTTTCAAGGATTGGGTTGTCATTAACGATATTGTGAATGCAGTGGAGCAG<br>CGTTTAAGCGAGGCCGCTTAG<br><br>Amino Acid Sequence:<br>MGSSHHHHHHSSGLVPRGSHMTNETVKAIYTTLAEYLDMPVSELKENTNLENELLLDST<br>ELVCIMVALEKSMDISLKNVPFKDWVVINDIVNAVEQRLSEAA |

**Table S2**. Thermal stability curve fit of vulcPPT.

| Curve Fitting Model | Logistic |
|---|---|
| Equation | y = A2 + (A1-A2)/(1 + (x/x0)^p) |
| A1 | -8033.27984 ± 40.53121 |
| A2 | -962.00728 ± 37.40994 |
| x0 | 45.52884 ± 0.14873 |
| E50 | 45.52884 ± 0.14873 |
| p | 14.08977 ± 0.56409 |
| Reduced Chi-Sqr | 21690.66673 |
| R-Square (COD) | 0.99812 |
| Adj. R-Square | 0.99797 |

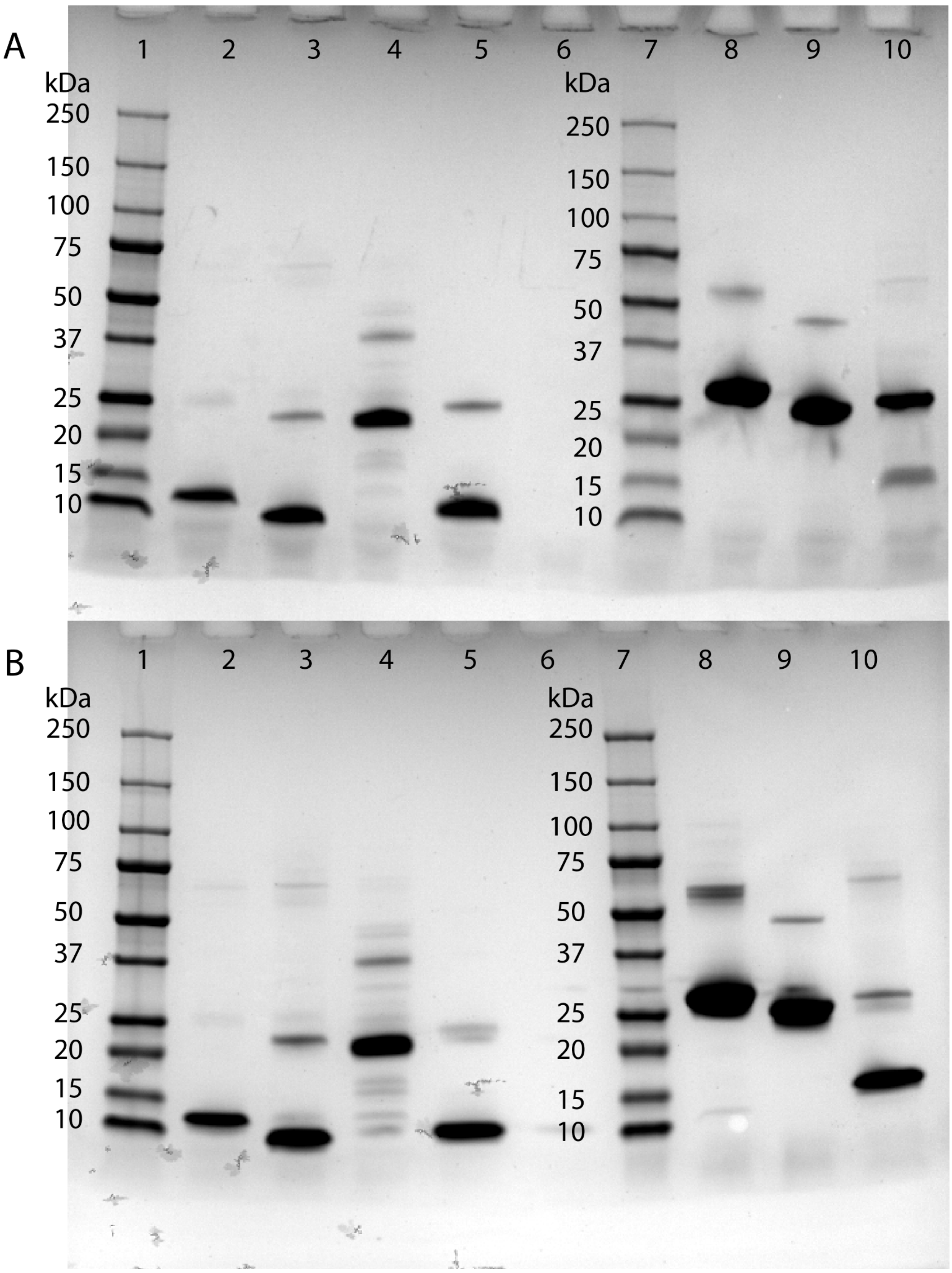


**Figure S1**. SDS-PAGE of the expressed and purified acyl carrier proteins and phosphopantetheinyl transferases used in this study. Gels were run under (A) non-reducing conditions and (B) reducing conditions with 5% (v/v) β-mercaptoethanol. In both gels, lanes 1–10 represent: (1+7) Precision Plus Protein™ Standards All Blue protein ladder (Bio-Rad); (2) vulcACP (12.1 kDa); (3) actACP (11.4 kDa); (4) AcpP (10.8 kDa); (5) zooACP (11.4 kDa); (6) *empty*; (8) vulcPPT (30.7 kDa); (9) Sfp R4-4 (27.3 kDa); (10) AcpS (16.2 kDa).

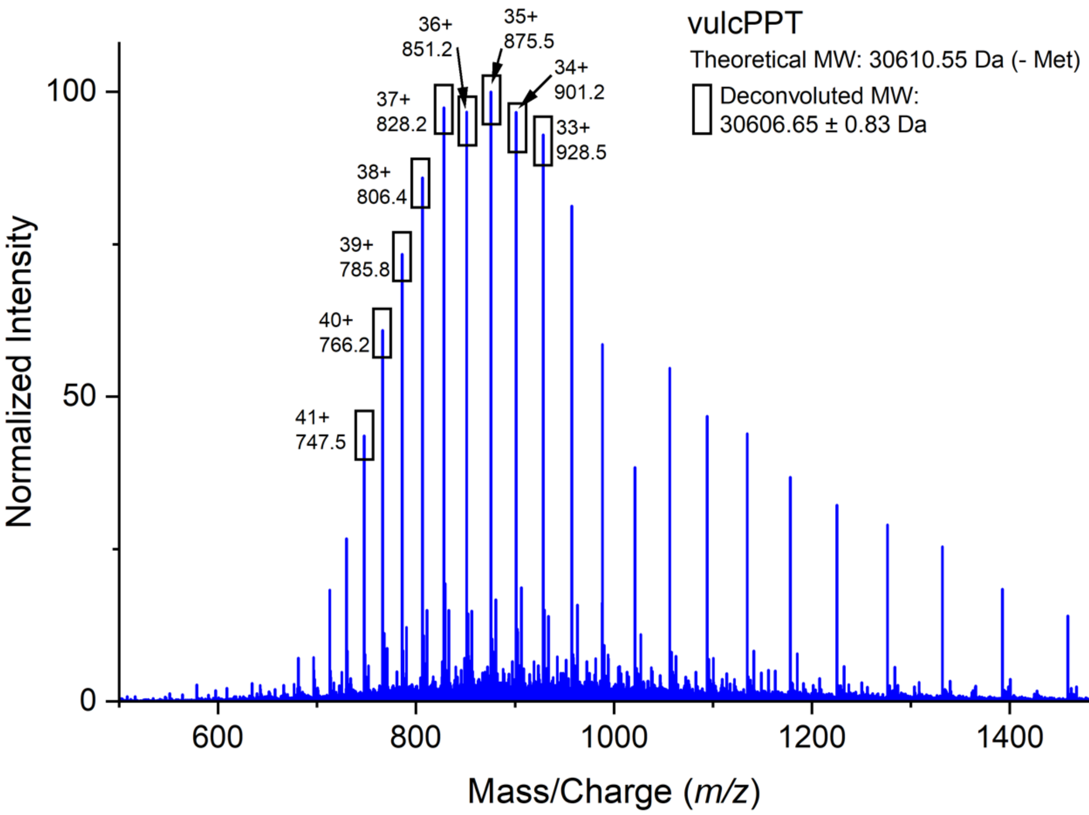


**Figure S2.** LC-MS spectrum of purified vulcPPT. The first nine major peaks were used for molecular weight deconvolution.

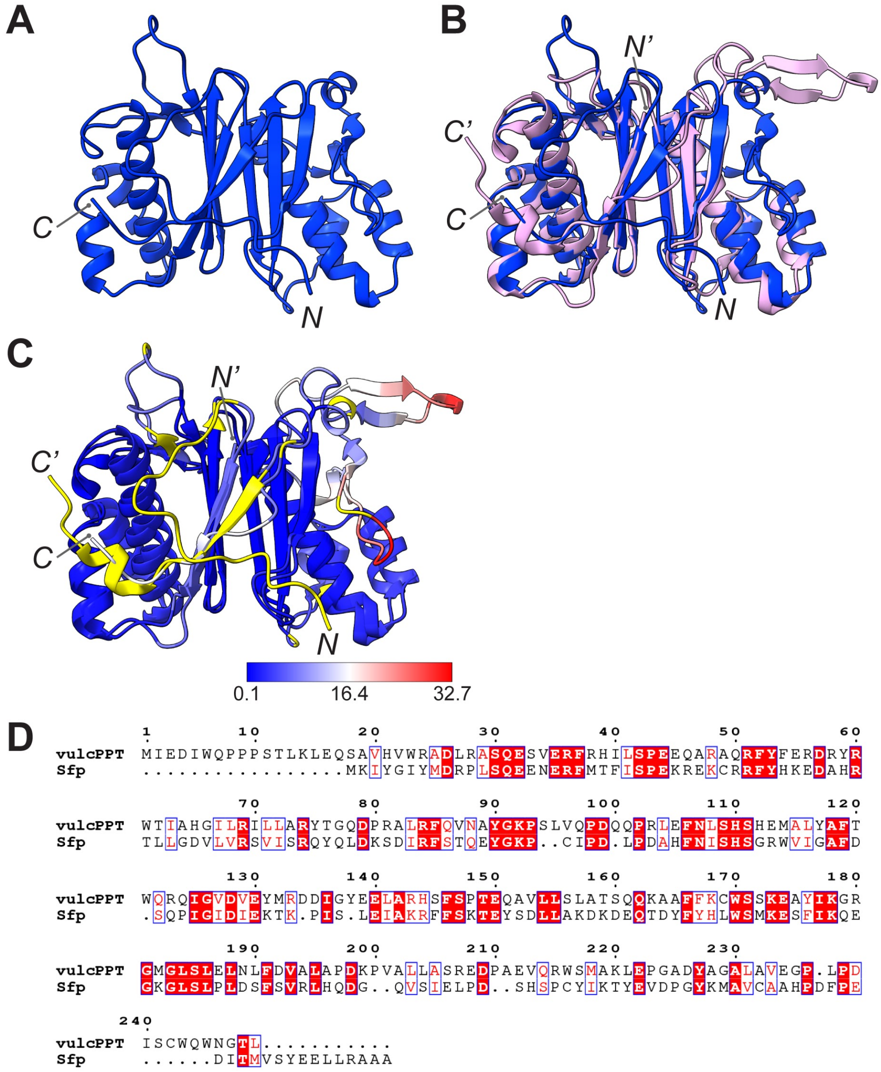


**Figure S3.** Structural comparison of vulcPPT and Sfp highlights their high structural similarity. (A) Predicted structure of vulcPPT generated using AlphaFold 3,[6] with the vulcPPT protein insert as input (see Table S1). The helicity was calculated to be 35.3%. (B) Structural overlay with Sfp (PDB 1QR0, pink) emphasizes their similarity. (C) The calculated RMSD between 127 pruned atom pairs is 1.089 Å. The color key indicates the sequence RMSD range from 0.1 Å (blue) to 32.7 Å (red), with unmatched residues shown in yellow. The *N*- and *C*-termini are labeled as *N* and *C* for vulcPPT, and *N'* and *C'* for Sfp. (D) Sequence alignment, performed using Clustal Omega[7] and ESPript 3,[2] indicates residue similarities as follows: red box with white character for strict identity, red character for group similarity, blue frame for similarity across groups, and black character for no similarity. The sequence from PDB 1QR0 (wild-type Sfp) was used for the alignment, while the R4-4 mutant of Sfp (Sfp R4-4)[8] was utilized in the experimental assays. The R4-4 mutant contains specific amino acid substitutions that enhance its substrate promiscuity compared to the wild-type enzyme: Lys28 (AAA) → Glu (GAG), Lys31 (AAA) → Lys (AAG), Thr44 (ACC) → Glu (GAG), His90 (CAC) → His (CAT), and Lys136 (AAG) → Lys (AAG).

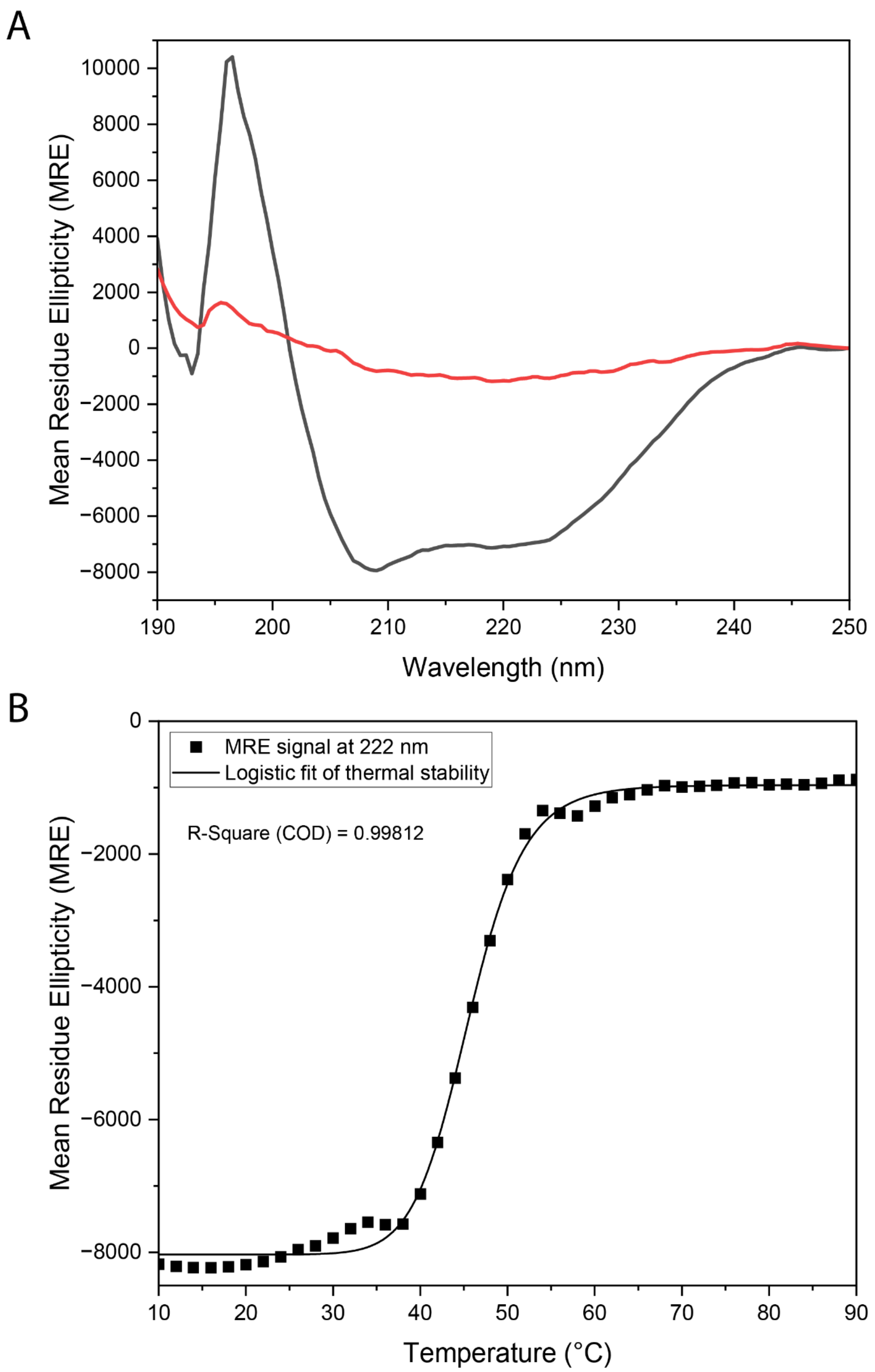


**Figure S4.** Circular dichroism spectrum and $T_{melt}$ of vulcPPT. (A) Dark line represents the CD signal of folded vulcPPT in the far-UV range (190–250 nm) at 25 °C, prior to collecting the $T_{melt}$ thermal stability curve of vulcPPT. Red line represents the CD signal in the far-UV range of denatured vulcPPT after completing the $T_{melt}$ thermal stability curve of vulcPPT and cooling back down to 10 °C. VulcPPT does not refold after denaturation. (B) $T_{melt}$ thermal stability curve of vulcPPT. CD signal at 222 nm was collected over a temperature range of 10–90 °C in 2 °C steps. All spectra were blank-corrected and a sigmoidal fit using a logistic function was fitted onto the data. E50/x0 value from the sigmoidal fit corresponded to a $T_{melt}$ of 45.53 ± 0.149 °C. A second transition between 30–35 °C is possible, which could suggest a multi-domain structure or a unit of secondary structure that is partially stabilized through contact with the structure of the rest of the protein.

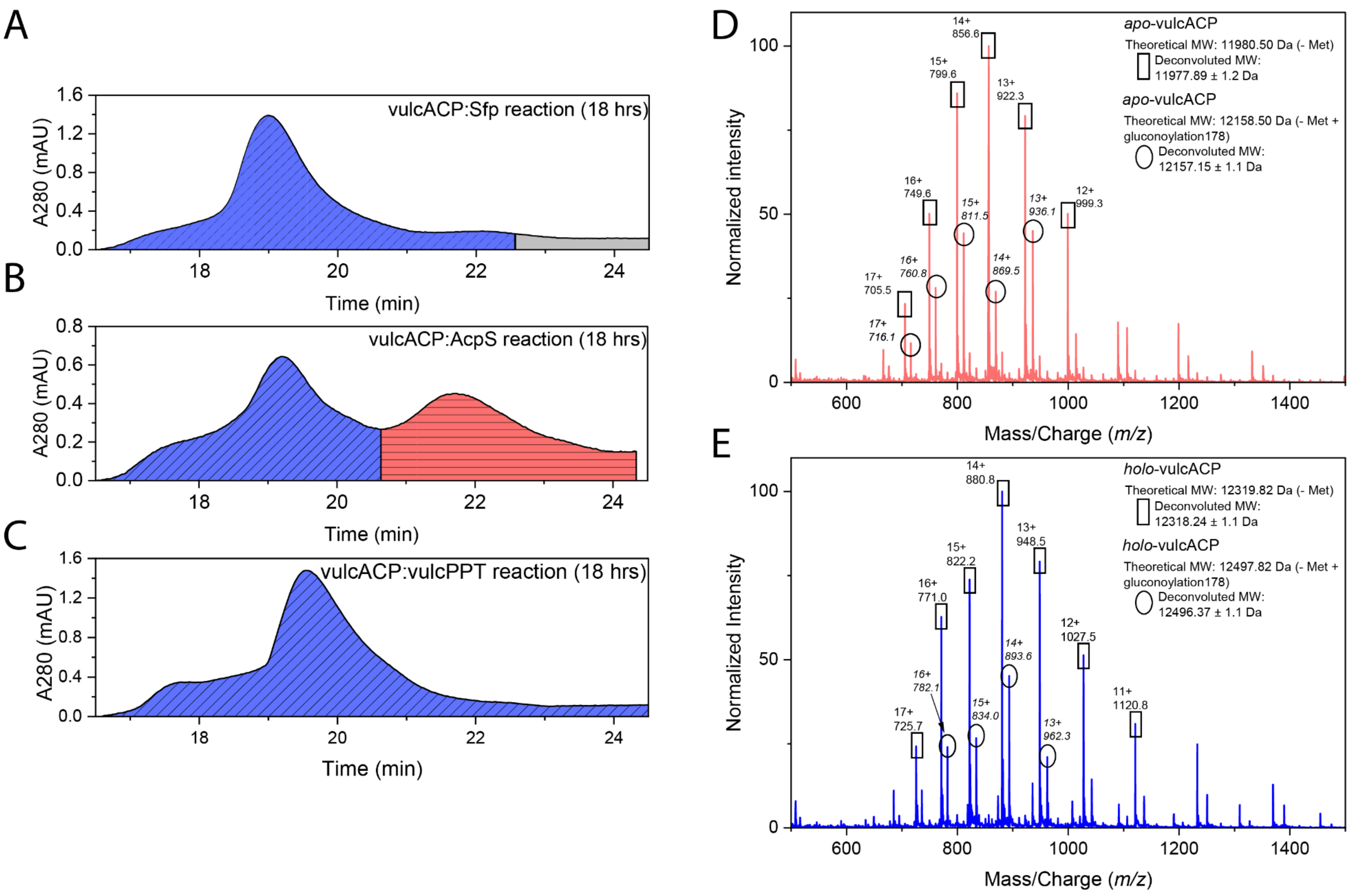


**Figure S5.** Mass spectra of *apo*- and *holo*-vulcACP, and LC chromatograms after PPTases reactions with Sfp, AcpS and vulcPPT. *Apo*- (red) and *holo*- (blue) ACP peaks were confirmed by mass-spectrometry. Areas without visible ACP are denoted in gray. (A) UV-Vis spectrum of the reaction between vulcACP and Sfp, showing full conversion to the *holo* form. (B) UV-Vis spectrum of the reaction between vulcACP and AcpS, showing an incomplete conversion. As a result, a mixture of *holo* and *apo* is observed. (C) UV-Vis spectrum of the reaction between vulcACP and vulcPPT, showing full conversion to the *holo* form. (D) Representative deconvoluted mass spectrum of *apo*-vulcACP. (E) Representative deconvoluted mass spectrum of *holo*-vulcACP.

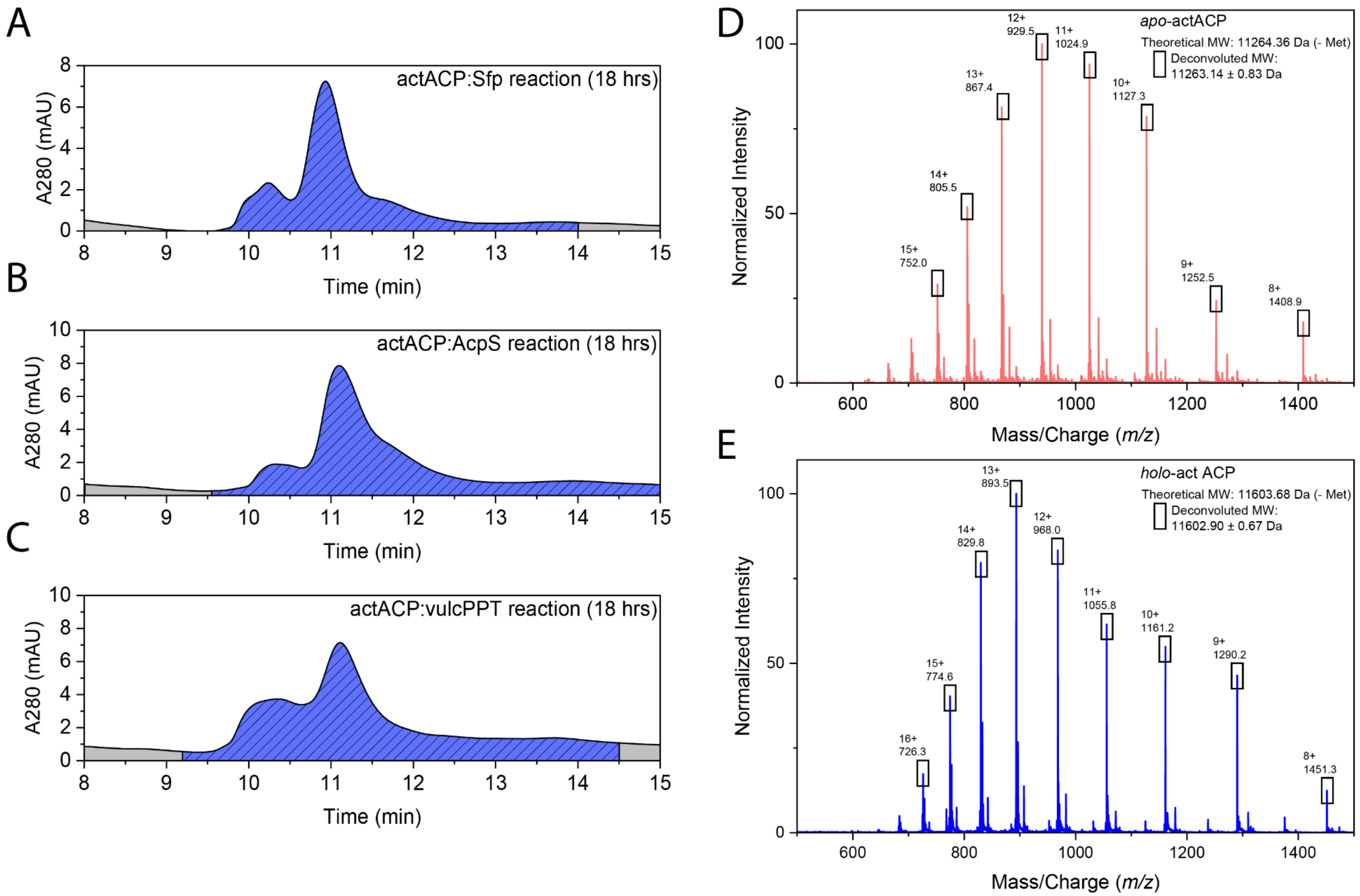


**Figure S6**. Mass spectra of *apo*- and *holo*-actACP, and LC chromatograms after PPTases reactions with Sfp, AcpS and vulcPPT. *Apo*- (red) and *holo*- (blue) ACP peaks were confirmed by mass-spectrometry. Areas without visible ACP are denoted in gray. (A) UV-Vis spectrum of the reaction between actACP and Sfp. (B) UV-Vis spectrum of the reaction between actACP and AcpS. (C) UV-Vis spectrum of the reaction between actACP and vulcPPT. All three PPTases show full conversion to the *holo* form. (D) Representative deconvoluted mass spectrum of *apo*-actACP. (E) Representative deconvoluted mass spectrum of *holo*-actACP.

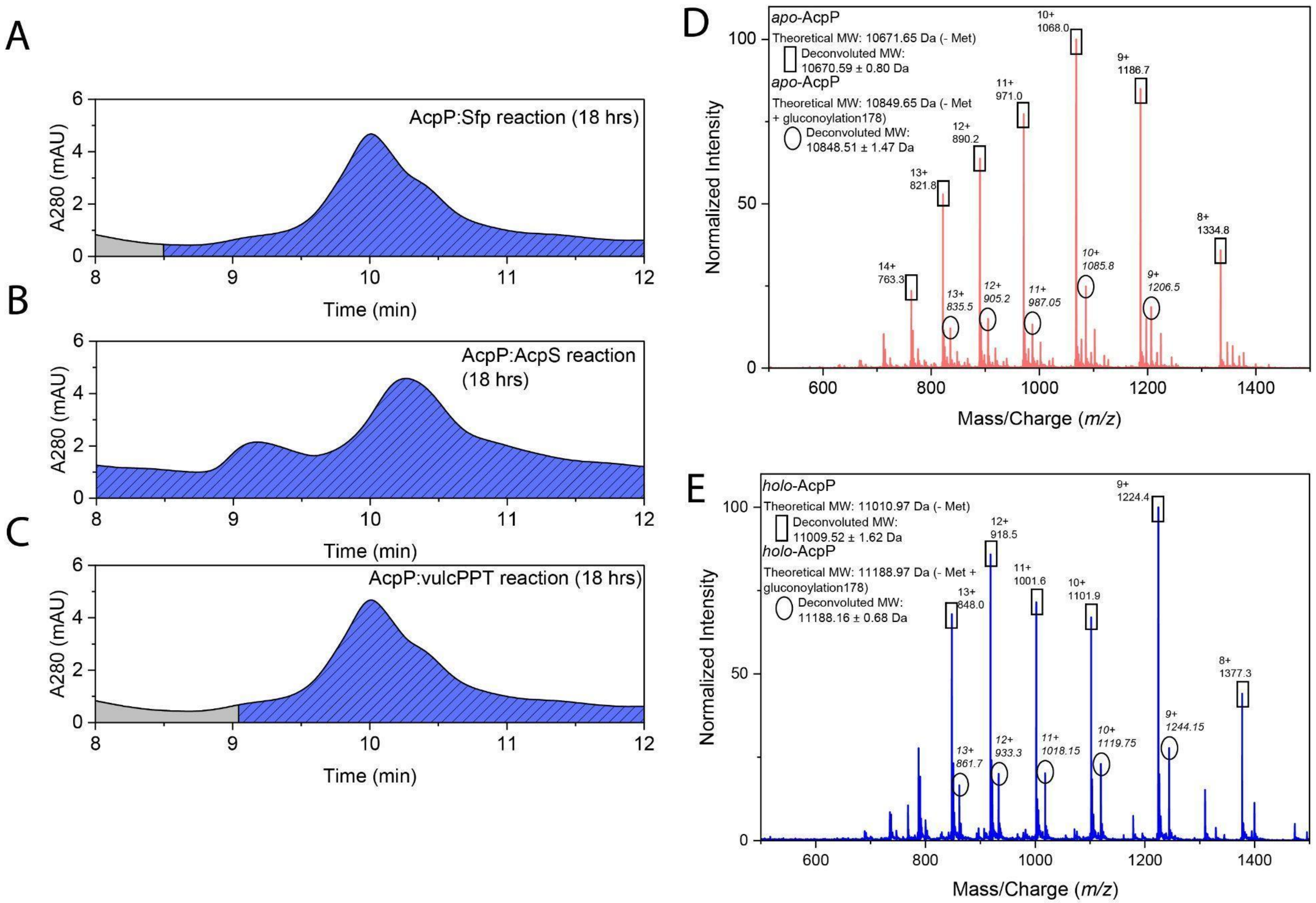


**Figure S7**. Mass spectra of *apo*- and *holo*-AcpP, and LC chromatograms after PPTases reactions with Sfp, AcpS and vulcPPT. *Apo*- (red) and *holo*- (blue) ACP peaks were confirmed by mass-spectrometry. Areas without visible ACP are denoted in gray. (A) UV-Vis spectrum of the reaction between AcpP and Sfp. (B) UV-Vis spectrum of the reaction between AcpP and AcpS. (C) UV-Vis spectrum of the reaction between AcpP and vulcPPT. All three PPTases show full conversion to the *holo* form. (D) Representative deconvoluted mass spectrum of *apo*-AcpP. (E) Representative deconvoluted mass spectrum of *holo*-AcpP.

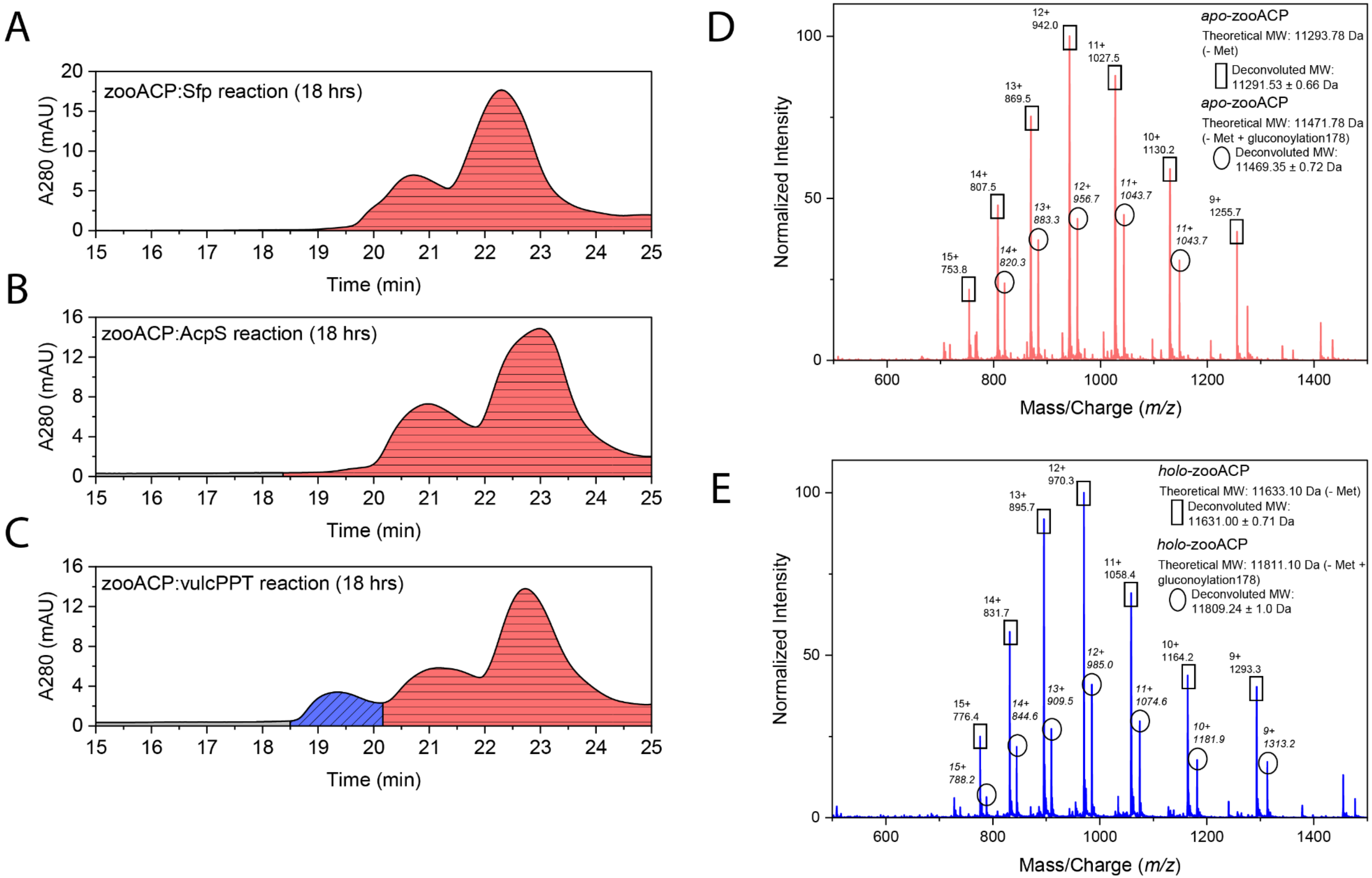


**Figure S8**. Mass spectra of *apo*- and *holo*-zooACP, and LC chromatograms after PPTases reactions with Sfp, AcpS and vulcPPT. *Apo*- (red) and *holo*- (blue) ACP peaks were confirmed by mass-spectrometry. Areas without visible ACP are denoted in gray. (A) UV-Vis spectrum of the reaction between zooACP and Sfp, showing no conversion to the *holo* form. (B) UV-Vis spectrum of the reaction between zooACP and AcpS, showing no conversion to the *holo* form. (C) UV-Vis spectrum of the reaction between zooACP and vulcPPT, showing an incomplete conversion to the *holo* form. As a result, a mixture of *holo* and *apo* is observed. (D) Representative deconvoluted mass spectrum of *apo*-zooACP. (E) Representative deconvoluted mass spectrum of *holo*-zooACP.

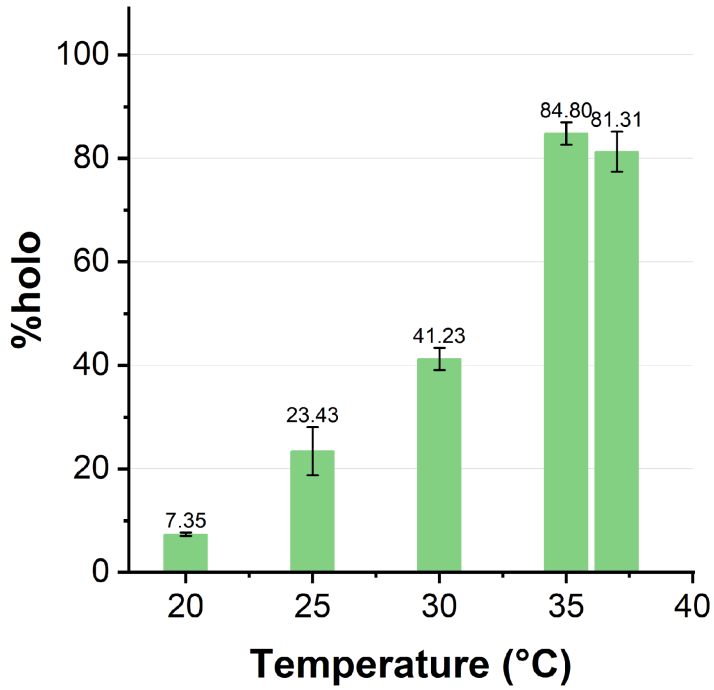


**Figure S9**. Temperature optimization of the phosphopantetheinylation of *apo*-zooACP by vulcPPT (detailed version of Fig 3A) at pH 7.6, demonstrating that 35 °C results in the highest conversion (84%) to the *holo* form.

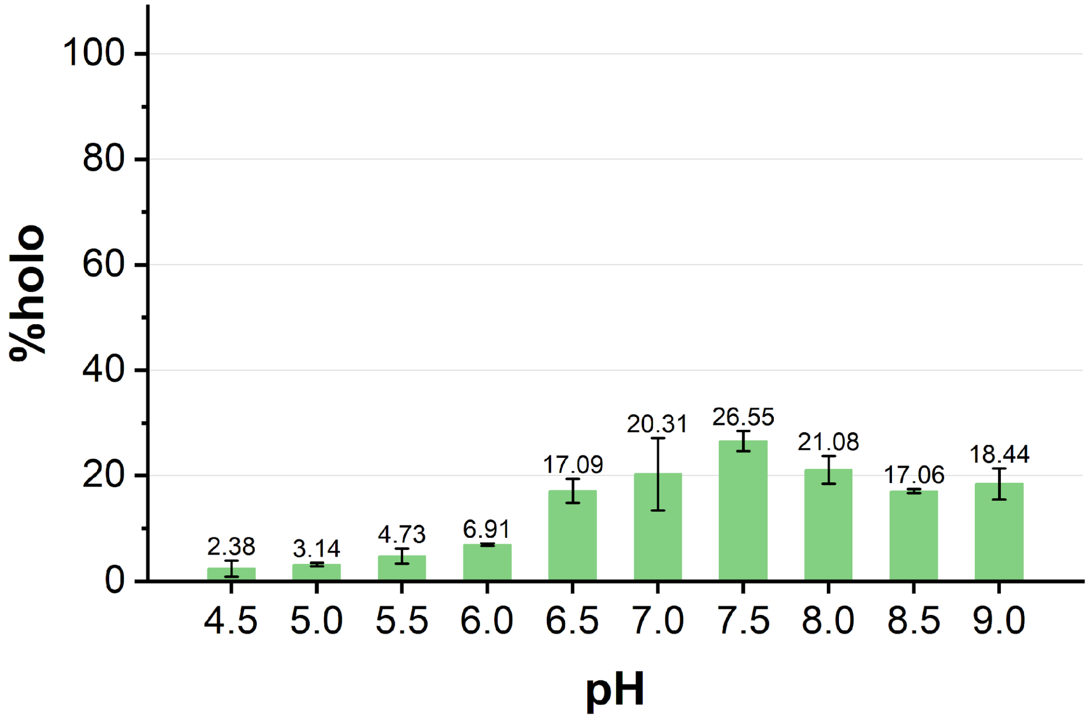


**Figure S10**. pH optimization of the phosphopantetheinylation of *apo*-zooACP by vulcPPT (detailed version of Fig 3B) at 25 °C, demonstrating that pH 7.5 results in the highest conversion (27%) to the *holo* form.

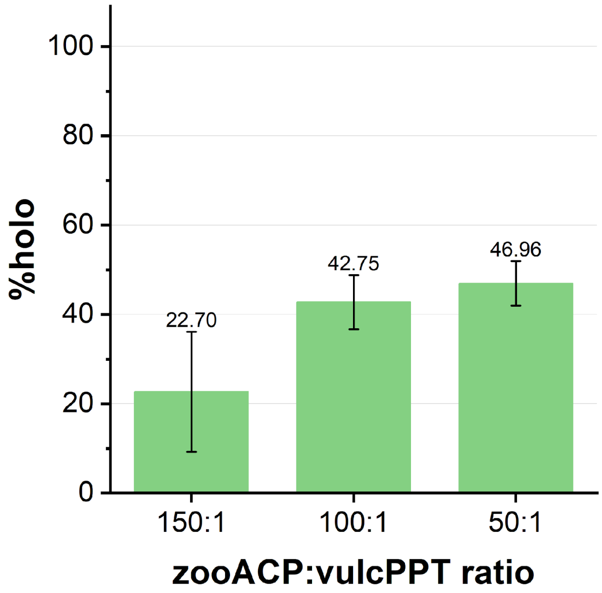


**Figure S11**. VulcPPT concentration optimization of the phosphopantetheinylation of *apo*-zooACP by vulcPPT. Three ratios of differing molar concentrations were tested: 150:1 (150 µM ACP to 1.0 µM vulcPPT), 100:1 (150 µM ACP to 1.5 µM vulcPPT) and 50:1 (150 µM ACP to 3.0 µM vulcPPT). The 50:1 zooACP:vulcPPT ratio showed the highest conversion to the *holo* form (47%), although within the standard deviation of the 100:1 ratio.

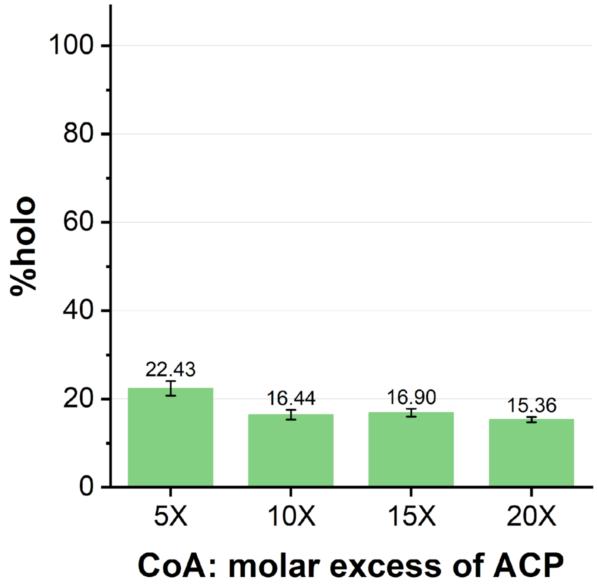


**Figure S12**. Coenzyme A concentration optimization of the phosphopantetheinylation of *apo*-zooACP by vulcPPT. Four CoA concentrations were tested at molar excesses of 5x (750 µM), 10x (1.50 mM), 15x (2.25 mM), and 20x (3.00 mM) relative to 150 µM *apo*-zooACP. The 5x CoA concentration resulted in the highest conversion to the *holo* form (22%), though the significance was not major compared to the other CoA concentrations.

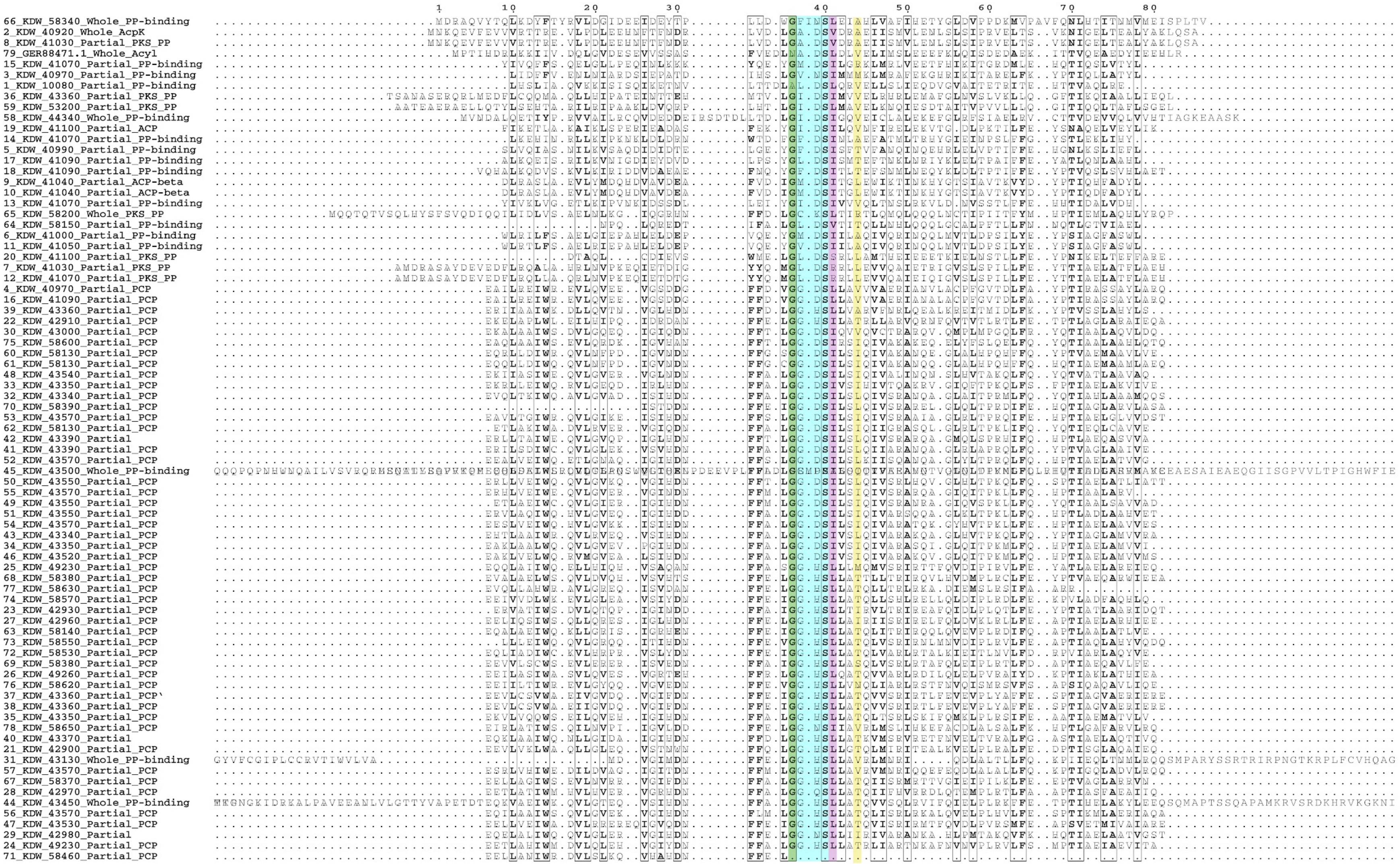

**Figure S13. Multiple sequence alignment of 79 carrier proteins encoded by the *Dictyobacter vulcani* sp. W12 genome.** 97.5% (77/79) of the CPs have a G as the first residue of the motif. Only 6.3% of the CPs shared the traditional Sfp-favored sequence “DSL”, instead having DSI (44.3%, 35/79), HSL (34.1%, 27/79), or some other amino acid triad in its place.
